# A deep learning approach for improving two-photon vascular imaging speeds

**DOI:** 10.1101/2022.11.30.518528

**Authors:** Annie Zhou, Samuel A. Mihelic, Shaun A. Engelmann, Alankrit Tomar, Andrew K. Dunn, Vagheesh M. Narasimhan

## Abstract

A potential method for tracking neurovascular disease progression over time in preclinical models is multiphoton fluorescence microscopy (MPM), which can image cerebral vasculature with capillary-level resolution. However, obtaining high-quality, three-dimensional images with a traditional point scanning MPM is time-consuming and limits sample sizes for chronic studies. Here, we present a convolutional neural network-based algorithm for fast upscaling of low-resolution or sparsely sampled images and combine it with a segmentation-less vectorization process for 3D reconstruction and statistical analysis of vascular network structure. In doing so, we also demonstrate that the use of semi-synthetic training data can replace the expensive and arduous process of acquiring low- and high-resolution training pairs without compromising vectorization outcomes, and thus open the possibility of utilizing such approaches for other MPM tasks where collecting training data is challenging. We applied our approach to large field of view images and show that our method generalizes across imaging depths, disease states and other differences in neurovasculature. Our pre-trained models and lightweight architecture can be used to reduce MPM imaging time by up to fourfold without any changes in underlying hardware, thereby enabling deployability across a range of settings.

## Introduction

The neurovascular network transports chemicals (e.g., oxygen, nutrients, waste) to and from the brain to support neuronal activity. Neurovascular function is disrupted by disorders such as stroke, Alzheimer’s and other neurodegenerative diseases, and diabetes, with lasting effects that are not fully understood^1,2^. Advances in multiphoton fluorescence microscopy (MPM) have enabled imaging with capillary-level resolution *in vivo*, and this noninvasive tool could be used to monitor capillary-level changes over time in cerebral vasculature as a potential predictor of disease progression/prognosis^3–6^. A constraint with MPM, however, is the slow acquisition process that is necessary for producing high quality, three-dimensional images with a traditional point scanning multiphoton imaging setup. Given the physical limitations of a live animal, the acquisition speed puts a limit on study sizes and the ability to reach statistically significant results. Although previous approaches have sought to improve imaging speeds by incorporating innovative imaging hardware, these implementations come at high cost and complexity and cannot be readily employed to existing infrastructure^7–10^.

An alternative, more cost-effective and accessible approach might be to computationally improve the image acquisition process using Convolutional Neural Networks (CNNs), which leverage existing datasets of MPM images. While several recent advances have been made in applying CNNs to improve MPM imaging results^11–16^, to our knowledge, only one has been focused on improving MPM imaging speed. Guan et al. presented a CNN for improving the imaging speed of a two-photon fiberscope for neuronal imaging using a conditional generative adversarial network (cGAN)^16^. They achieved a 10-fold speedup in frame rate. A drawback to their approach, however, is the requirement for a two-part training set, involving both *ex vivo* and *in vivo* imaging, which is extremely expensive and time-consuming. Several other models for general denoising or segmentation for MPM have also been focused primarily on neuronal or calcium imaging^11–14^, with only one to our knowledge focused on vascular segmentation^15^–none of which is used for improving vascular imaging speeds.

Here, we describe a CNN based approach trained to take images captured at low-resolution (128 × 128 pixels), thereby at much faster speeds, and then upscale these to a higher resolution (512 × 512 pixels), without compromising the accuracy of vascular morphological information that is extracted or introducing additional noise. The upscaling process from low- to high-resolution using deep learning is referred to as image super-resolution. We then combined this with a vectorization pipeline to obtain quantitative statistics of neurovasculature. Our pre-trained models and light architecture allows for fast acquisition, image super-resolution, and vectorization of MPM images without the limitations of added hardware and can be used to reduce imaging time by up to fourfold.

## Results

### Structure and analysis pipeline overview

Our pipeline to improve two-photon microscopy acquisition and vectorization accepts individual as well as multiple frames from MPM imaging. Our process is split into two main parts: an image super-resolution CNN designed to upscale low-resolution images and a vectorization pipeline that is designed to output morphological statistics on the acquired images (**Fig. 1**). For super-resolving the images, we used the PSSR Res-U-Net architecture, which had been shown to restore images of presynaptic vesicles and neuronal mitochondria from a scanning electron microscope (SEM) and a laser scanning confocal microscope, respectively^17^. We utilized this architecture over several other possible ones because it: (a) allowed us to use semi-synthetic data for training which circumvents the need to acquire real-world image pairs for training, which is difficult and expensive for large datasets (b) enabled us to also employ multi-frame inputs that could leverage information across correlated images at similar depths (c) does not utilize an adversarial network in the training, which are more challenging to train as well as to evaluate the generated models (d) allowed us to use a transfer learning approach to initialize our model with weights obtained with the architecture trained on ImageNet, a large natural image classification dataset. For vectorization, we used Segmentation-Less, Automated, Vascular Vectorization (SLAVV)^18^, which uses simple models of vascular anatomy, and efficient linear filtering and vector extraction algorithms with manual or automated vector classification. Using a multi-frame PSSR approach and combining it with a vectorization pipeline, we show that we are able to restore two-photon vascular images sufficiently for extracting morphological characteristics through vectorization.

**Fig. 1:**
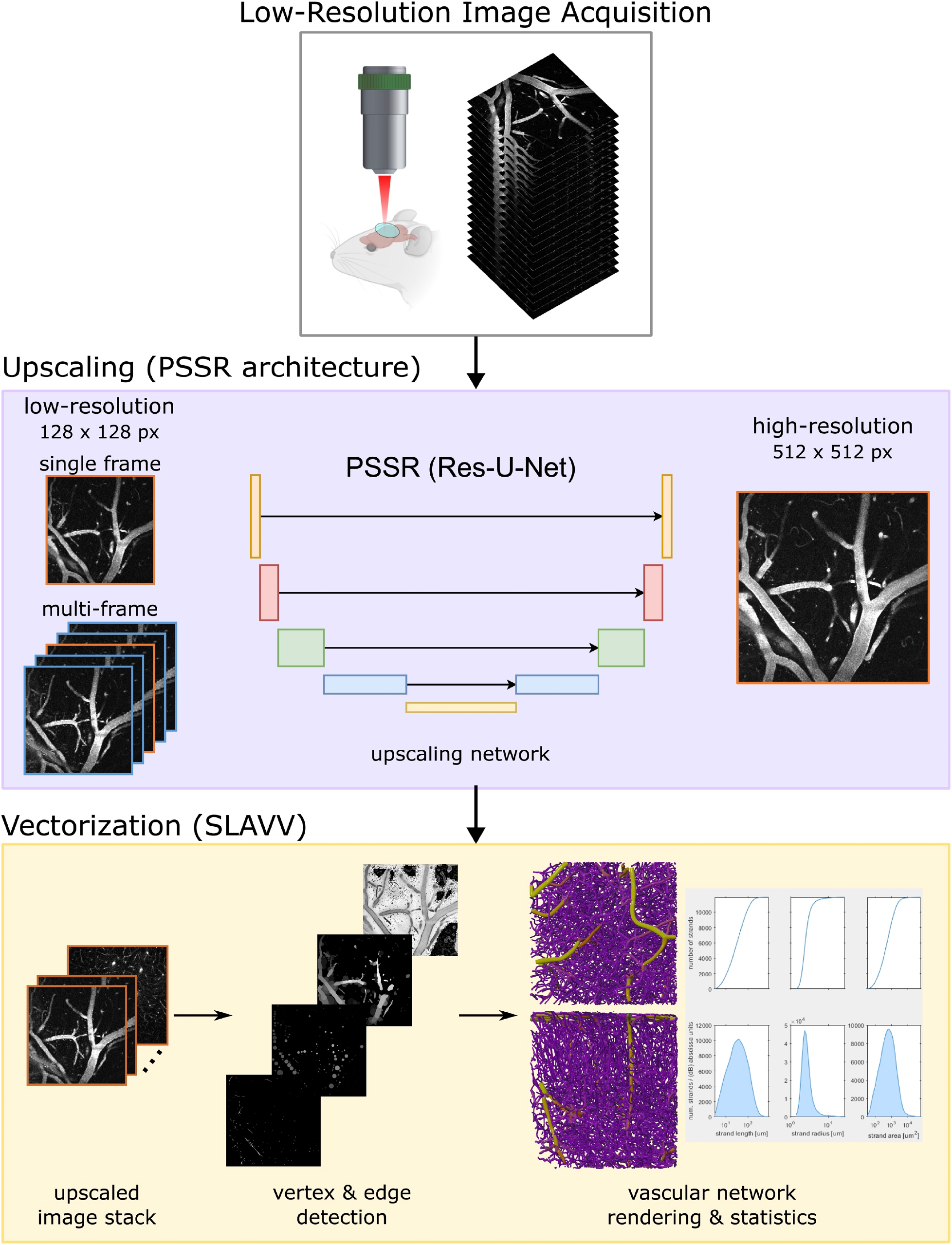
Structure and analysis pipeline. Low-resolution images (128 × 128 pixels) are acquired using two-photon microscopy. A deep learning (PSSR Res-U-Net) based upscaling process generates high-resolution images (512 × 512 pixels), which take much longer to acquire, from low-resolution images. Segmentation-less vascular vectorization (SLAVV) generates 3D renderings and calculates network statistics from an upscaled image stack.

### Transfer learning, creation and evaluation of semi-synthetic training data

Traditional approaches to upscaling images involve acquiring paired high- and low-resolution real-world images that we could use for training the model. For our task, however, this process is expensive and time-consuming, and in certain situations impossible for live animals. This challenge greatly limits the practical sample size of training datasets. To overcome this difficulty, we sought to use semi-synthetic training data that mimics low-resolution acquisition to greatly improve sample size. Semi-synthetic training data was created by adding noise to, then downscaling, full-resolution images from a two-photon vascular image repository of previously acquired images (see Data Availability). We evaluated several approaches for the creation of this semi-synthetic data and compared our results to a model trained with a real-world dataset of the same sample size.

To mimic the noise observed in real-acquired low-resolution images (i.e., real data), we tested models trained with semi-synthetic images that were created with the following noise filters: no noise (downscaled only, used as the reference), Poisson noise, Gaussian noise, and additive-Gaussian-distributed noise (**Fig. 2a**). Real-acquired low-resolution images served as input images to test the model. We evaluated model performance using standard image quality metrics, specifically, by calculating peak signal-to-noise ratio (PSNR) and structural similarity index measure (SSIM) between the model output and acquired full resolution image.

**Fig. 2:**
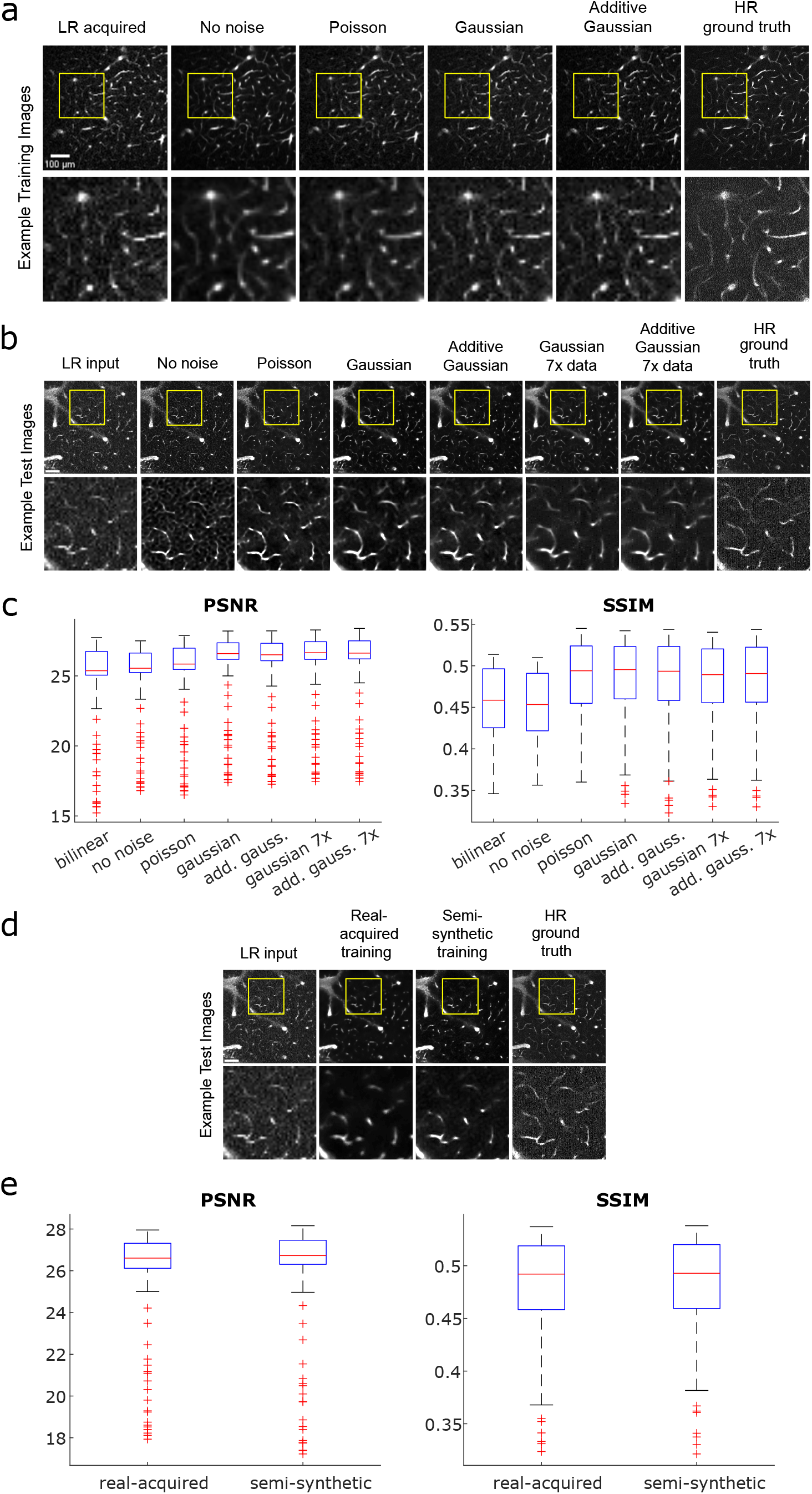
Generating and evaluating semi-synthetic training data. **a** Examples of semi-synthetic training images created using different types of added noise prior to downscaling: no noise (downscaling only), Poisson, Gaussian, and additive Gaussian. Acquired low-resolution (LR, 128×128 pixels) and high-resolution (HR, 512×512 pixels) ground truth images are shown for reference. **b** Resulting test images from models trained using each noise method, with acquired low-resolution image for model input and acquired high-resolution image as ground truth for comparison. All models were trained with 3,399 image pairs, with the Gaussian and additive Gaussian models further tested on 24,069 image pairs (7x) to further test performance. **c** Boxplot comparison of PSNR and SSIM values for each noise method image in **b** measured against ground truth image. Values plotted for an image stack of 222 images. **d** Comparison of test images from models trained using real-world acquired vs. semi-synthetic data, with real-acquired low-resolution image for model input and acquired high-resolution image as ground truth for reference. All models were trained with 234 image pairs, a large reduction from the noise model comparison, due to the limited availability of real-world pairs. **e** Boxplot comparison of PSNR and SSIM values for real-acquired vs. semi-synthetic model outputs corresponding to **d**, measured against high-resolution ground truth image. All values are plotted for an image stack of 222 images.

The resulting median PSNR and SSIM values from each model, ranked from highest to lowest for both metrics, were as follows: Gaussian, additive-Gaussian, Poisson, no noise (**Fig. 2b-c**). This was determined using Wilcoxon signed rank tests with P < 0.005 (Bonferroni-adjusted). We noticed that the Gaussian and additive-Gaussian models perform similarly, and thus performed further testing to compare the two noise methods using a larger training set, consisting of 24,069 semi-synthetic training image pairs—7x the preliminary training set of 3,399 semi-synthetic image pairs. The test image outputs from the models trained with the larger dataset showed notable qualitative improvements, with fewer false detections, less noise, and smoother vessel shapes. With the larger dataset, the Gaussian model produced a slightly higher median PSNR value but did not produce a median SSIM value that was statistically significantly different from that produced by additive-Gaussian (Wilcoxon signed rank test, P < 0.005). Despite the slightly higher PSNR performance by the Gaussian model, however, a qualitative comparison suggested that the additive-Gaussian results had somewhat less noise and higher sensitivity to fainter vessels. Additionally, PSNR only measures similarity in pixel values and does not necessarily predict vectorization performance, which is what we ultimately wish to optimize. To fully validate and compare the performance between the Gaussian- and additive-Gaussian-trained models, we performed a final comparison test using the Segmentation-Less Automated Vascular Vectorization (SLAVV) software (further described in later section). We found that the additive-Gaussian model output allowed for a more accurate vessel detection overall compared to the Gaussian model output (95.7% vs 95.6%). Based on these results, we chose to perform subsequent analyses using the model trained with the full dataset of semi-synthetic images created using the additive-Gaussian noise method to maximize accurate vessel detection.

To evaluate the effectiveness of using semi-synthetic data in place of real-world training data, we compared the performance of models trained with each method (**Fig. 2d-e**). For this comparison, both models were trained with 234 image pairs, due to the limited availability of acquired image pairs. The output image from the real-acquired model appeared blurrier and over-predicted vessel diameters more significantly compared to the semi-synthetic model (**Supplemental Figure 1**). Nonetheless, the model trained with real-world data had higher median PSNR and SSIM values compared to the model trained with semi-synthetic data (Wilcoxon signed rank test, P < 0.05), although the values were close (PSNR: 26.9 vs. 26.6; SSIM: 0.492 vs. 0.494). We deem the results similar enough for semi-synthetic training data to be used in place of real-world training data. The use of semi-synthetic data advantageously circumvents complications from imprecise alignment in the acquisition of image pairs, limited availability of existing images (677 pairs), and high material and labor costs for data collection.

### Single-frame vs. multi-frame training

A key issue with low-resolution acquired images is the diminished amount of total signal capture. This can result in noisier images and cause spurious vessels to appear in the vectorization process. A potential method for reducing false detections is providing the model neighboring depth images on a stack, which are highly correlated in signal but not in noise. Thus, we sought to improve the performance of our model by using multi-frame image input.

We compare the performance of the single-frame model to a multi-frame model, with an additional comparison against the traditional bilinear upscaling method, for both semi-synthetic and real-world test images (**Fig. 3**). The traditional bilinear upscaling method offers a baseline performance measure for a non-CNN approach. The multi-frame model is trained with input image stacks consisting of five sequential images in depth, with axial offsets of 0.3 μm, to predict a single output image—the third image in the input sequence. The multi-frame model yields images with higher PSNR and SSIM values than the single-frame model, and both PSSR methods outperform the bilinear upscaling method for both semi-synthetic and real-world test images (Wilcoxon signed rank test, P < 0.0167, Bonferroni-adjusted). Overall PSNR and SSIM values are higher for semi-synthetic images compared to real-world images, which is unsurprising given the model was trained completely on semi-synthetic images. Nonetheless, the real-world output images from our models show that individual vessels can be resolved, which is much more important for the final vectorization process than the exact pixel values measured by PSNR.

**Fig. 3:**
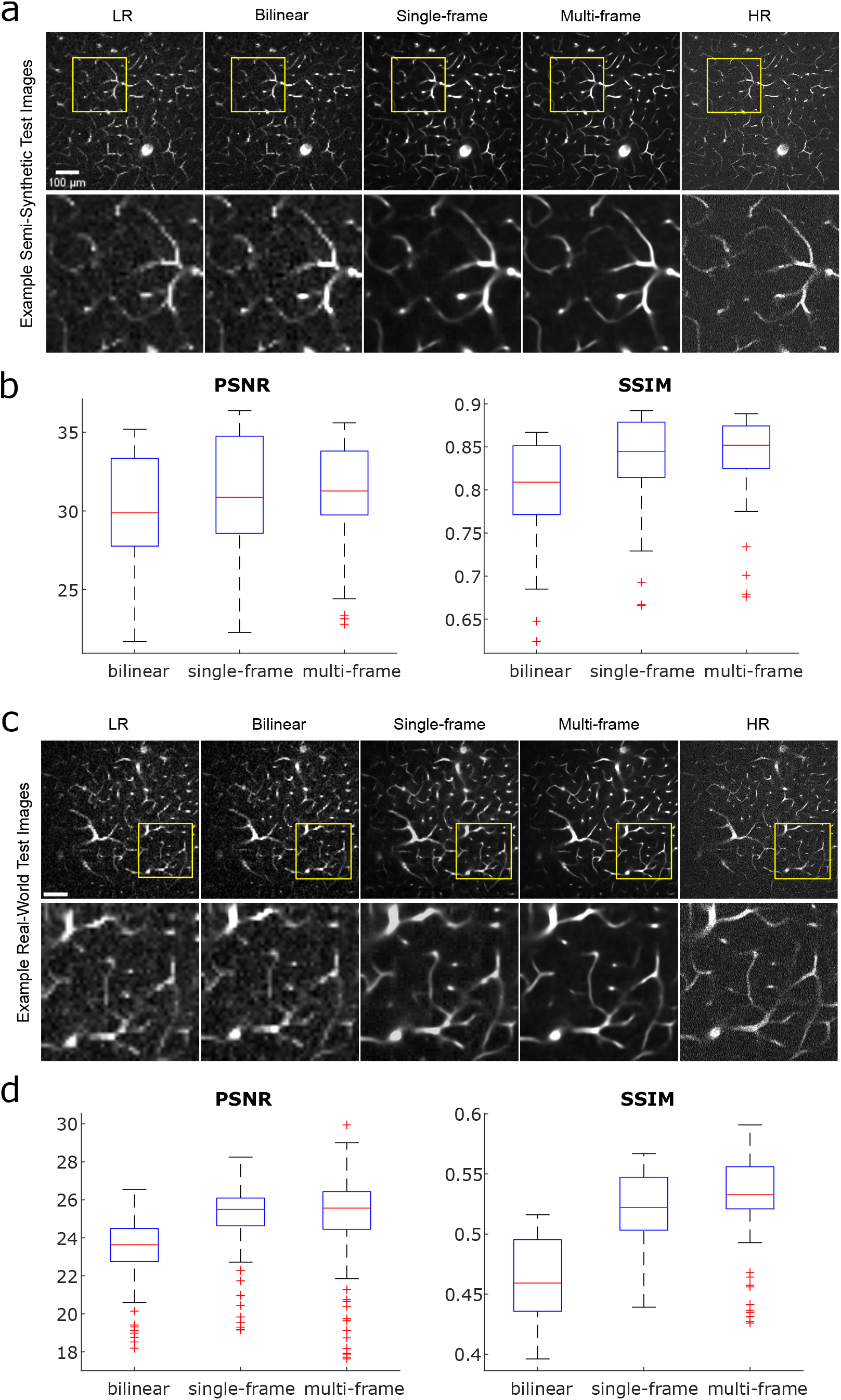
Comparison of performance between bilinear upscaling, a single-frame model, and a multi-frame model for semi-synthetic and real-acquired test images. All models were trained with 24,069 image pairs. **a** Semi-synthetic test images from bilinear upscaling and models trained using single-vs. multi-frame data. Acquired low-resolution image for model input and acquired high-resolution image as ground truth are shown for reference. **b** PSNR and SSIM plots corresponding to semi-synthetic test results from **a**. **c** Real-world test images from bilinear upscaling and models trained using single-vs. multi-frame data. **d** PSNR and SSIM plots corresponding to real-world test image results from **c**.

### Reconstruction and stitching of infarct images

A major application of two-photon imaging that we aim to make more accessible with our approach is large field of view (FOV) imaging of diseased vasculature. An example of this is acquiring images from a stroke model, which is of interest for studying disease effects on vascular morphology. Large FOV imaging with high resolution is a time-consuming process and thus would benefit substantially from the speedup offered by low-resolution imaging. Large FOV images are achieved by acquiring, then stitching standard-sized tiles together using ImageJ’s Grid/Collection Stitching plugin^19^. However, the long acquisition times required of current technologies limit our ability to collect substantial two-photon image sets of diseased vascular networks and thus limit the availability of images of diseased vasculature that could be used for training data.

To investigate the feasibility of using our models to drastically reduce imaging times for large FOV images of diseased vasculature with minimal information loss, we examined the ability of our single-frame and multi-frame models, trained only with images of normal vasculature, to restore a semi-synthetic large FOV image of an ischemic infarct (four weeks post-stroke) collected in a preliminary study (**Fig. 4**). The differences in morphology between vasculature in the peri-infarct region and normal vasculature are exemplified by the differences between the top half of the full image, which more closely resembles normal vasculature, and the bottom half of the image, which captures the infarct region and the more immediately surrounding vessels. Ischemic infarct vessels appear significantly more parallel to the imaging plane, thus creating image slices with higher vascular area density compared to the more perpendicularly oriented vessels further from the infarct. Despite these morphological differences and having only trained with images of normal vasculature, our models are able to resolve capillaries in the infarct region, as shown in the insets of **Fig. 4**. The multi-frame output image more closely resembles the HR image compared to the single-frame image, as vessel radii are more consistent in the multi-frame image. In the case of the LR and bilinear-upscaled images, the individual capillaries in the inset cannot be resolved.

**Fig. 4.**
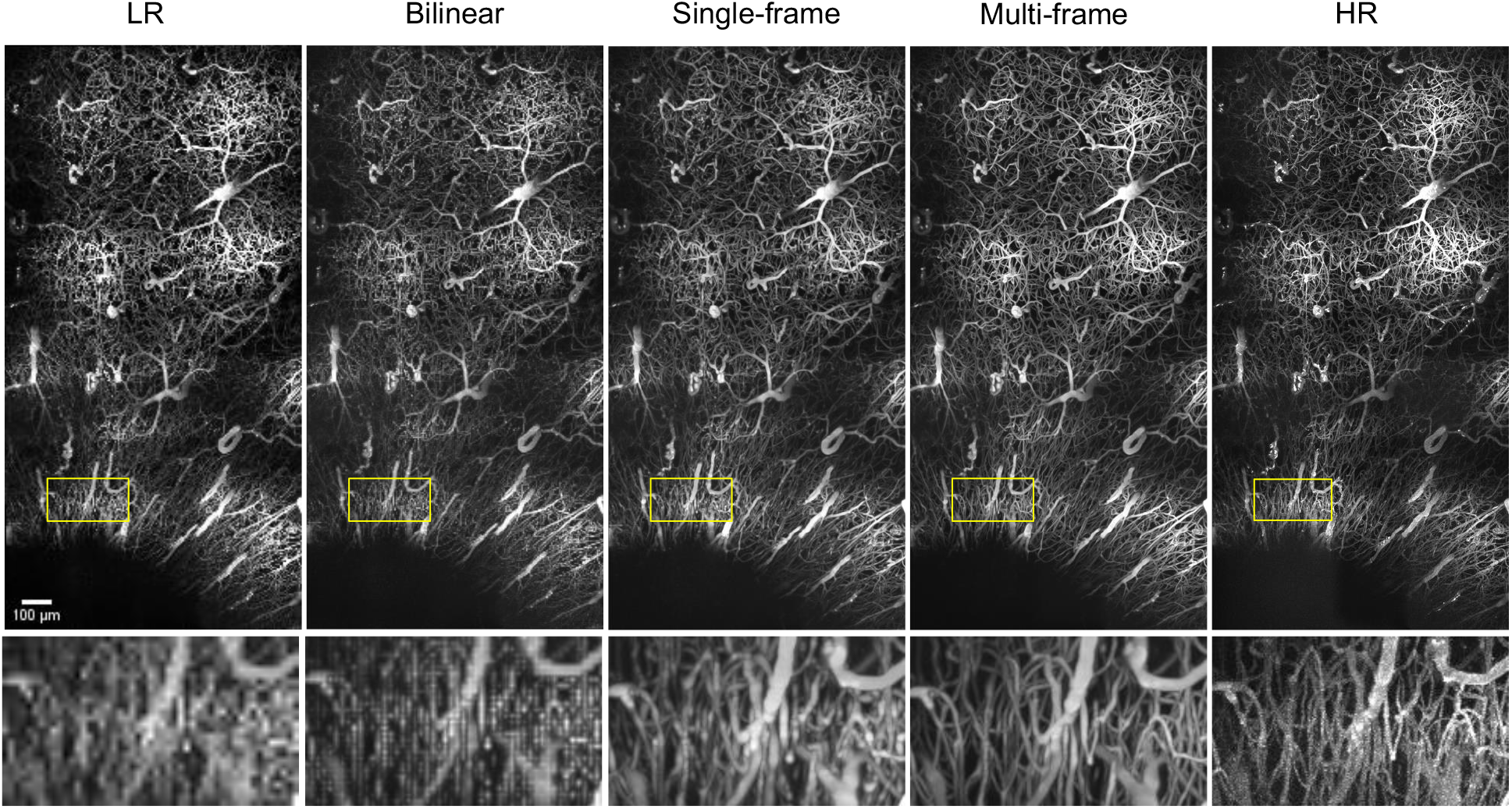
Maximum intensity projections (x-y) of ischemic infarct images consisting of 2×4 tiles with 213 slices (final dimensions 1.18 mm × 2.10 mm × 0.636 mm, pixel dimensions 1.34 μm × 1.36 μm × 3 μm) for a semi-synthetic low-resolution image, bilinear upscaled image, single- and multi-frame output images, and acquired high-resolution image.

### Vectorization

Vectorization is the ultimate step that extracts quantitative information for evaluating the vascular morphology of a network. Therefore, we are interested in comparing different image generation strategies by comparing performance after vectorization. We demonstrate successful vectorization of single- and multi-frame model output images from real-acquired low-resolution images using manual-curation-assisted SLAVV and visualization with VessMorphoVis^20^ (**Fig. 5a**). Additionally, we perform a more objective comparison of our models’ performance using a previously described method^18^, which uses simulated images from a known ground truth and automated vector classification (no manual assist). Using a known ground truth (derived from the real-acquired high resolution vectorized network shown in (**Fig. 5a**), we are able to quantify the sensitivity, specificity, and accuracy of the vectorized upscaled images. We generated the simulated vascular image to have the same contrast-to-noise ratio (CNR) of 0.94 as the real acquired high resolution image and to be representative of the image quality of a typical image acquired by our two-photon microscope. We created a low-resolution version of the simulated image using the same method for creating semi-synthetic training data and then upscaled it using bilinear interpolation and our single-frame and multi-frame models. We then vectorized these images using fully automated (globally thresholded) SLAVV at peak segmentation performance (measured against the ground truth image). The resulting strand objects are the minimal set of one-dimensional traces which span the entire vascular network.

**Fig. 5.**
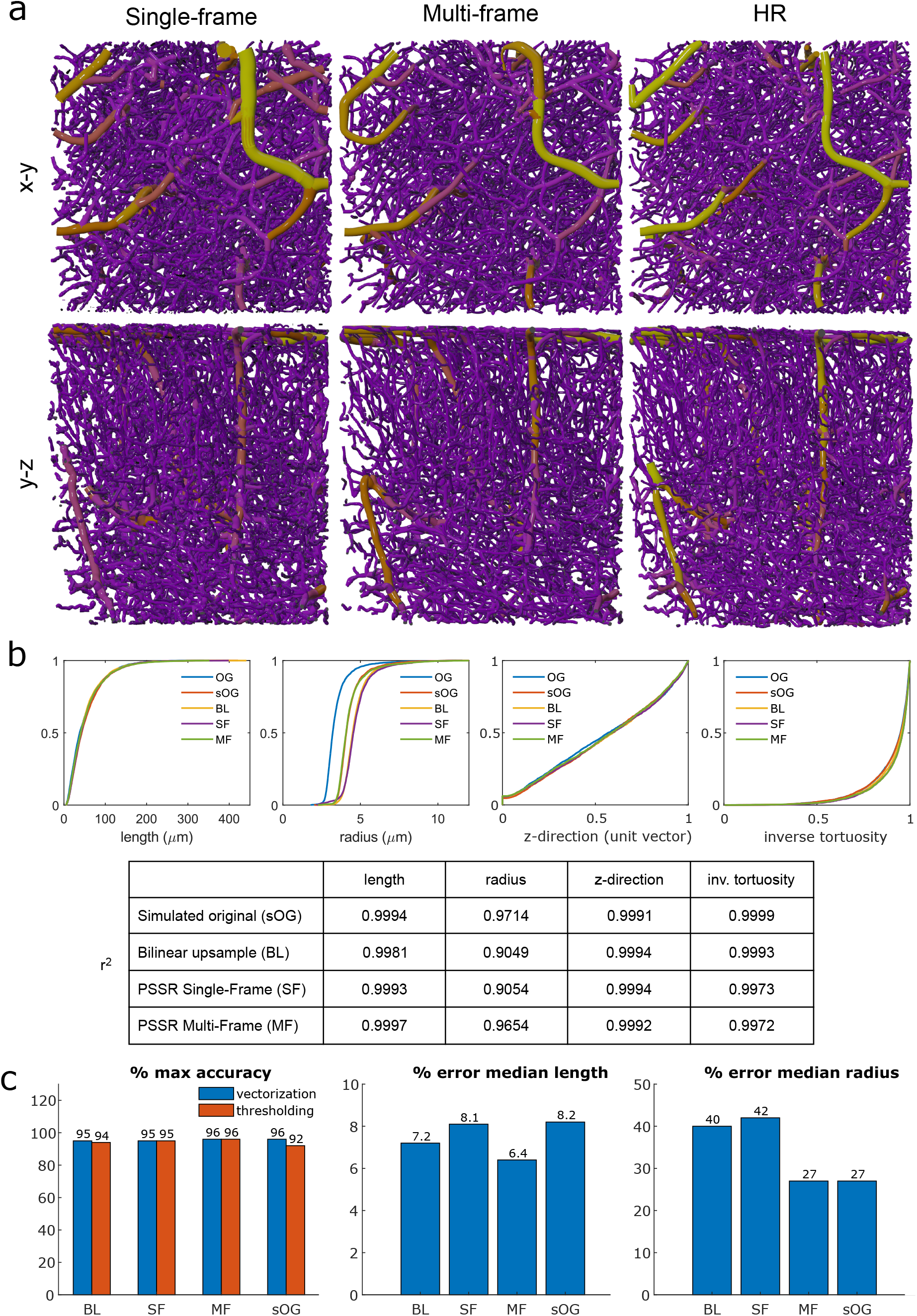
Vectorization results. **a** Blender rendering of vectorized images using VessMorphoVis^20^ for visual comparison between single- and multi-frame results and an acquired high-resolution image. We performed manual curation for this vectorization process. **b** Vectorized image statistics for automated curation process with known ground truth (simulated from manually curated high-resolution image). CDFs shown for metrics of length, radius, z-direction, and inverse tortuosity for original (OG), simulated original (sOG), bilinear upscaled (BL), and PSSR single- and multi-frame (SF, MF, respectively) images. Pearson’s correlation values (*r^2^*) were calculated between the original image and each simulated or upscaled image for each metric. **c** Statistics regarding maximum accuracy (%) achieved with vectorization or thresholding and % error in median length and radius for each method.

We plotted cumulative distribution functions (CDFs) for each image for each of the following strand metrics: length, radius, z-direction, and inverse tortuosity (**Fig. 5b**). We included the simulated original image in the analysis as a control for the automated curation process since the ground truth image was obtained through manual curation. For each strand metric, we calculated Pearson’s correlation (*r^2^*) values between the CDFs of the ground truth image and the simulated images. Of all the images, the simulated original image maintained the highest *r^2^* value for average strand radius and inverse tortuosity. Our multi-frame model had the highest *r^2^* value for strand length, while bilinear and the single-frame model produced the highest *r^2^* value for z-direction. We performed a Kolmogorov–Smirnov (K-S) test to compare the CDFs of each upscaling method against that of the simulated original image. The multi-frame CDFs for strand length, radius, and z-direction; the single-frame CDFs for length and z-direction; and the bilinear CDF for z-direction were not significantly different from those of the simulated original image (p < 0.0167, Bonferroni-adjusted). Thus, of the tested upscaling methods, the multi-frame model produced the most statistically comparable strand metrics to the simulated original image.

We calculated overall accuracy with respect to the ground truth for each image (**Fig. 5c**). The original simulated image retains the highest vectorization accuracy (96.2%), followed by multi-frame (95.7%), single-frame (95.2%), and bilinear (94.5%). In terms of accuracy with raw image segmentation through intensity thresholding, however, multi-frame performs best (96.0%), followed by single-frame (95.3%), bilinear (94.4%), and original simulated image (91.9%). We also calculated the percent error in the median length and median radius, the characteristics that best represent the vessel morphology, between each image against the ground truth values. Multi-frame produced the lowest median length error (6.4%), followed by bilinear (7.2%), single-frame (8.1%), and the original simulated image (8.2%). The bilinear and single-frame images had notably higher median radius errors (40.1% and 41.6%, respectively) compared to the multi-frame and original simulated images (both 26.9%), which was noted with visual inspection of the images as well. These statistics further support that a multi-frame upscaled image produces comparable vectorization results to an original high-resolution image.

## Discussion

To our knowledge, this is the first time that a deep learning model has been demonstrated to improve imaging speeds for two-photon microscopy by upscaling and denoising low-resolution images of vasculature while retaining accuracy in extracted morphological characteristics. For this application, our model outperforms the traditional, non-CNN bilinear upscaling technique in output image quality (**Fig. 3, 4**) and vectorization accuracy (**Fig. 5**). The performance of our models also improves notably with increased training data (**Fig. 2b**), therefore substantial time and material costs are reduced by training the model with semi-synthetic images generated from our database of 28,563 previously acquired two-photon vascular images. Real-world data also introduces further complications by requiring image registration. Not only would this add computational hours, but the image registration process also does not produce perfect alignment because it is limited to being purely translational and free of interpolation to maintain the original recorded pixel values. Any rotational misalignments would not be accurately correctable. We speculate that these misalignments in real-world training image pairs caused the overly blurry and enlarged vascular structures seen in the results of preliminary experiments (**Fig. 2d**, **Supplementary Fig. 1**).

Models trained from semi-synthetic images proved capable of restoring low-resolution vascular images and outperformed the standard bilinear interpolation method. For performing segmentation via intensity thresholding (**Fig. 5c**), multi-frame had the highest accuracy of the upscaling methods and significantly outperformed the original standard-resolution image. We hypothesize that this is a result of the denoising that occurs in the PSSR process. Since all upscaled images had higher intensity thresholding accuracy compared to the original image, we further postulate that upscaled images have less noise overall because fewer pixels are physically captured—all pixels created during upscaling are interpolated from neighboring pixel values and thus free of noise from the image acquisition process^21^. For performing vectorization, however, the diminished noise does not offset accuracy losses from the upscaling process. We determined that the multi-frame model yields the highest accuracy of the three upscaling methods but did not outperform the original standard-resolution image. Nonetheless, the multi-frame image also produced the greatest number of CDFs for strand metrics that were not significantly different from those of a standard-resolution image. We consider these results from the multi-frame model to be within acceptable tolerance for vectorization accuracy and similarity in strand statistics for our previously described purposes of characterizing the structural properties of the vessels of a particular network^22,23^.

By acquiring low-resolution images, the imaging time could be reduced by up to half in a two-photon microscope with a resonant-galvo scanning system and by up to fourfold with a galvo-galvo scanning system. The reduction in imaging time can have several benefits. For instance, faster imaging times reduce risk of phototoxicity and thermal damage if excitation powers are kept constant^24,25^. Additionally, the amount of time for which the subject is under anesthesia is reduced, decreasing the risk of vascular dilation^26^ which can create skewed vectorization statistics. The reduction in imaging time can also decrease the injection volume and frequency of fluorescent dye, which alone can save up to hundreds of dollars in addition to eliminate the risk of sample misalignment caused by a re-injection during an imaging session. A specific but major benefit for those wishing to conduct chronic studies is that the faster acquisition times will allow for larger cohorts, which are currently constrained by the number of animals that can reasonably be imaged within each timepoint. This would yield more statistically significant sample sizes for studying and comparing healthy and diseased vasculatures over time.

With the potential for future disease studies in mind, we tested our model on data from a mouse that was given a stroke. We show that our model, despite having only been trained with images of healthy vasculature, can reasonably restore images taken from a peri-infarct region with sufficient resolution for the image stitching algorithm to successfully create a large FOV image. These results from semi-synthetic test data are promising for being able to apply the model broadly to different disease models, although further validation should be performed with real-world test data.

Another area of research that could benefit from increased imaging speeds is in the study of light propagation through the brain, for the development of noninvasive brain imaging devices. With accessibility to a larger database of large FOV two-photon images, light propagation models can be more thoroughly developed, tested, and refined^27^. As precise capillary capture is not necessary for these models, the use of even lower resolution and faster imaging could be further explored with PSSR. Although losses in accuracy in low-resolution images are inevitable due to less information being captured, experimentation with the accuracy and speedup tradeoff can be done to fit the tolerance of any application.

The potential for further speedup by reducing frame averaging could also be explored. Lower frame averaging leads to higher noise levels, which PSSR can be used to reduce. With higher noise, we would expect increased false positive and/or false negative detections, leading to an overall reduction in restoration accuracy. A potential method that can be explored to combat this effect would be to modify the loss function to increase the penalty for false negative detections with the tradeoff of accepting more noise in the image. Alternatively, to prioritize denoising over having high sensitivity, the loss function could be modified to penalize false positive detections more heavily.

## Methods

### In vivo imaging

#### Animal Preparation

Cranial window implants were prepared in C57 mice with dura intact, as previously described^28^. During imaging, mice were anesthetized with isoflurane and body temperature was maintained at 37.5 °C. Blood plasma was fluorescently labeled with dextran conjugated Texas Red (70kDa, D1830, Thermo Fisher) dissolved in saline (5% w/v). The dye was administered intravenously via retro-orbital injection (0.1 mL). All animal protocols were approved by The University of Texas at Austin Institutional Animal Care and Use Committee.

For stroke model mice in **Fig. 4**, photothrombotic ischemia was induced through retro-orbitally injecting rose bengal (0.15ml at 15 mg/ml) and irradiating a penetrating arteriole branching from the middle cerebral artery for 15 minutes. The laser source had a 532 nm wavelength, a 20mW average power, and was focused to a ~300 μm diameter spot size. Mice were anesthetized with isoflurane (1.5%, 0.6-0.8 LPM) and body temperature was maintained with a heating pad during the procedure. Pial anatomy was visualized using laser speckle contrast imaging to select which artery to target and to confirm occlusion.

#### Image Acquisition

All images were acquired using a custom-built two-photon microscope, previously described^21^. The excitation source was an ytterbium fiber amplifier with an output beam of 1050 nm wavelength, 120 fs pulse width, and 80 MHz repetition rate^29^. High-resolution images were 512×512 pixels and low-resolution images were 128×128 pixels, both with a field size of 700×700 μm. Image stacks were acquired with 3 μm axial spacing. A resonant-galvanometer scanning system was used^21^, with average pixel dwell times of 87.8 ns and 20-frame averaging. Power at the sample did not exceed 170 mW and was identical between low- and high-resolution pairs. Large field of view images were taken as a 2-by-4 grid of standard images, with ~25-30% overlap between tiles.

### Image processing

#### Image pre-processing

All images were normalized prior to use as a training image or semi-synthetic test image. A 3D median filter of size [1 1 1] was applied to raw image stacks, followed by a full-scale contrast stretch (FSCS) to fill the 16-bit range with 0.3% saturation across the entire stack, using the normalization function provided by Fiji ImageJ^30^. This FSCS normalization method was determined to create the best images compared to FSCS across the entire stack without saturation and FSCS by image slice (**Supplementary Fig. 2**). Images were then converted from 16-bit to 8-bit and separated into individual frames.

#### Stitching

Large field of view images acquired as a 2-by-4 grid of standard images were stitched together using ImageJ’s Grid/Collection Stitching plugin^19^.

### Semi-synthetic image generation

#### Single-frame images

To create semi-synthetic low-resolution images for training, pre-processed real-world images received one of the following types of noise: Poisson, Gaussian (μ = 0, σ = 0.1), additive-Gaussian (μ = 0, σ = 5), or no noise prior to fourfold downscaling (from 512 × 512 pixels to 128 × 128 pixels). For additive-Gaussian noise, the local variance was scaled by 0.001. A range of parameters (i.e., mean, standard deviation, local variance) were tested to identify values for optimal performance. Models were trained for each combination of parameters and given test images. Output images were inspected visually, and image quality metrics (PSNR, SSIM) were calculated.

#### Multi-frame images

Low-resolution images from the single-frame semi-synthetic image generation step were used to create multi-frame training images. Multi-frame images consisted of five low-resolution images sequential in axial space with 0.3 μm separation (axial distance between acquired images).

### Neural networks and training

The Res-U-Net architecture as described by Fang et al. as PSSR (point-scanning super resolution) was used for single-frame and multi-frame training^17^. We used an MSE loss function after determining that L1 and feature loss did not perform as well (**Supplementary Fig. 3**). A learning rate of 9e-4 was used for single-frame training and 1e-4 was used for multi-frame training.

#### Training/Test Images

Preliminary training for finding the best noise model was done with 3,399 training (data from 5 mice, 16 stacks, 6 imaging sessions) and 676 validation image pairs (2 mice, 3 stacks, 2 imaging sessions). Final full dataset training was completed using 24,069 training (6 mice, 114 stacks, 19 imaging sessions) and 4,494 validation (6 mice, 22 stacks, 7 imaging sessions) image pairs. Real-acquired image pairs (677 image pairs from 2 mice, 3 stacks, 2 imaging sessions) were used for testing and evaluating the models. These image pairs were also used for comparing the performance between training with real-acquired pairs vs. semi-synthetic pairs (234 image pairs for training, 221 image pairs for validation, 222 image pairs for testing; each set from 1 mouse, 1 stack, 1 imaging session).

#### Hardware

Training was performed using Frontera at the Texas Advanced Computing Center (TACC) with four NVIDIA Quadro RTX 5000 GPUs using the CUDA version 10.0 toolkit.

### Image Quality Evaluation

Image quality between upscaled images and the original image was preliminarily assessed with peak signal-to-noise ratio (PSNR) and structural similarity index measure (SSIM). Both metrics were computed using built-in MATLAB functions. In combination, these metrics gave a general sense for image similarity, but were not indicators for morphological accuracy from vectorization. Generally, higher PSNR and SSIM values are desired. However, both metrics correlate similarities in raw intensity values to higher similarity between images, whereas the vectorization process is designed for vascular networks, is the end-goal of image acquisition, and produces quantitative anatomical information which may or may not be of interest to a particular researcher. For simplicity, when possible, we chose image segmentation accuracy (with respect to the ground truth) to measure general vectorization performance. In the case of unknown ground truth, SLAVV was used to segment the original image and estimate the CNR, which was used to match the quality of the simulated and real-acquired images. PSNR and SSIM values do not seem to fully reflect image quality improvement from denoising, as seen in **Fig. 2b** where image quality improves visually with the increased training data, but PSNR and SSIM values decrease slightly. In addition, higher variations in predicted pixel intensity value are seen within white vessel regions, which can cause lower PSNR and SSIM values despite not affecting visual quality or vectorization performance. This especially affects images closer to the surface of the brain, where large arteries dominate the image, and accounts for the outlier points seen in the boxplots of **Fig. 2** and **Fig. 3**.

### Vectorization

All vectorization was performed using SLAVV software^18^. Real acquired low-resolution images were upscaled using PSSR, then vectorized and manually curated. To obtain an objective comparison of methods without manual curation bias, a simulated image with known ground truth was created from the manually curated high-resolution image. The simulated image had an identical CNR (0.94) to the original acquired high-resolution image, as measured by SLAVV as:

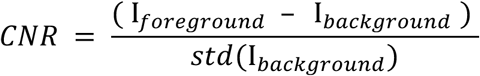

where I_foreground_ is average foreground intensity and I_background_ is average background intensity.

A low-resolution image was created using the previously described semi-synthetic image generation method. The low-resolution image was upscaled using bilinear interpolation, the single-frame model, and the multi-frame model. Vectorization of simulated images using automated curation was possible with the known ground truth, as previously described^18^.

From the vectorized networks for each upscaled image, the original simulated image, and the ground truth network, cumulative distribution functions were calculated for strand statistics (length, average radius, average z-direction, and inverse tortuosity). Two comparisons were then made for each strand statistics: Pearson’s correlation between each upscaled or original simulated image CDF and ground truth CDF; and Kolmogorov–Smirnov (K-S) test between each upscaled image and original simulated image. The Pearson’s correlation values compare the performance of each simulated (upscaled or original) image against the ground truth. The original simulated image serves as a baseline for vectorization performance. The K-S test is used to determine whether the performance of each model is significantly different compared to the original simulated image.

Overall accuracy was calculated for comparison as follows:

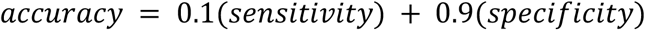

with *sensitivity* and *specificity* defined as follows:

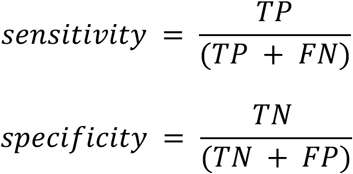

where TP is the number of true positive detections, TN is true negative, FP is false positive, and FN is false negative.

Blender renderings for vectorization visualization were created using VessMorphoVis software^20^.

### Statistical Analysis

All significance values were Bonferroni-adjusted from the standard P value of 0.05 to address the increased possibility of type-I error.

## Data availability

All data used for training, validation, and testing are available at the following link: (link to dataverse)

## Code availability

All the code presented in this work together with the trained network model are freely available at the following link: (link to gitHub page)

## Acknowledgments

V.M.N. was supported on a grant for human brain evolution by the Allen Discovery Center program, a Paul G. Allen Frontiers Group advised program of the Paul G. Allen Family Foundation as well as a fellowship from the Good Systems for Ethical AI at The University of Texas at Austin. This work was also supported by the National Institutes of Health (NS108484 to A.K.D; 3T32EB007507 and 5T32LM012414 to A.Z.). Objective and mouse images in Figure 1 were adapted from Biorender. GPU and computing support for the project was supported on Director’s discretionary fund at the Texas Advanced Computing Cluster.

## Contributions

A.Z. conceived of the study. A.Z., S.E., and A.T. collected data. A.Z., S.M., and V.M.N. analyzed the imaging data and built the pipeline. A.Z. and V.M.N. wrote the manuscript with input from all co-authors. V.M.N. and A.D. supervised the study.

## Competing interests

The authors declare no competing interests.

## Notes

### Competing Interest Statement

The authors have declared no competing interest.

