## Supplementary Fig. for "A deep learning approach for improving two-photon vascular imaging speeds"

### Supplemental Figures

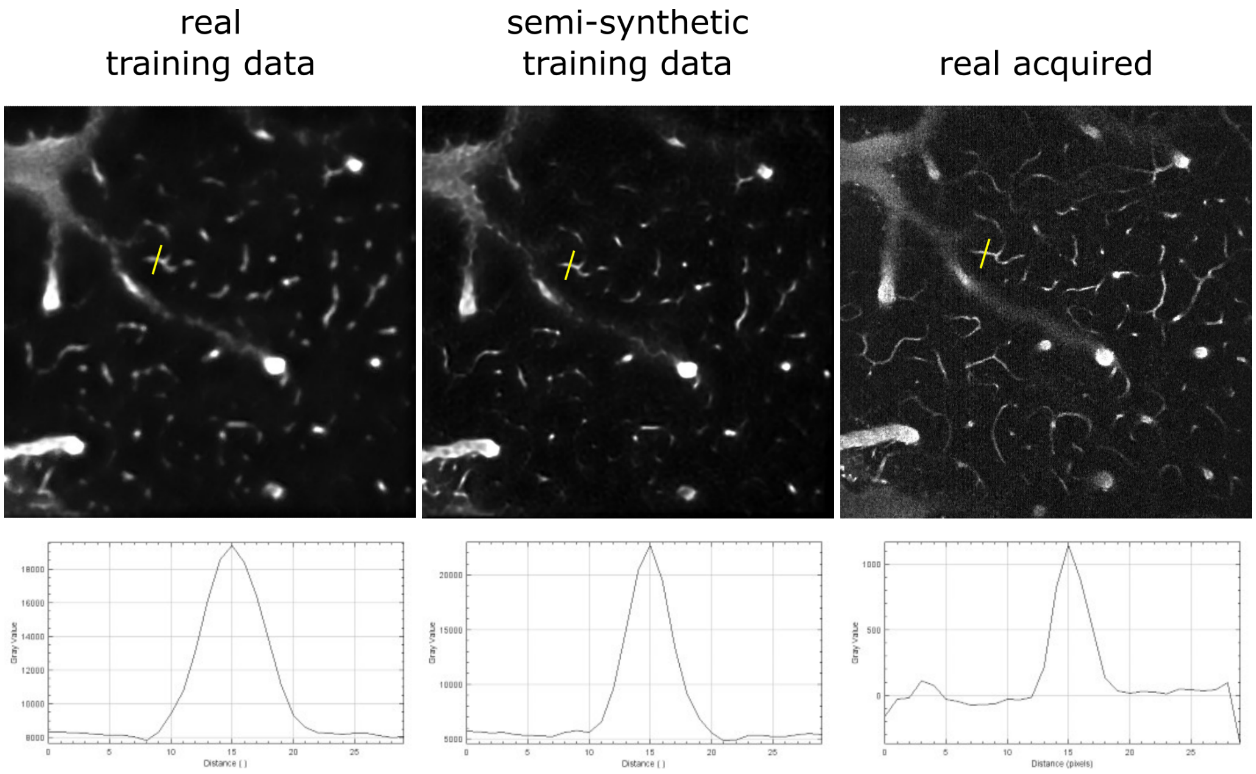


**S.Fig. 1. Vessel diameter comparison for test image outputs from models trained with real training data vs. semi-synthetic training data, vs. a real-acquired image.** Approximate vessel diameters are as follows in units of pixels: real training: 13, semi-synthetic training: 10, real acquired 7.

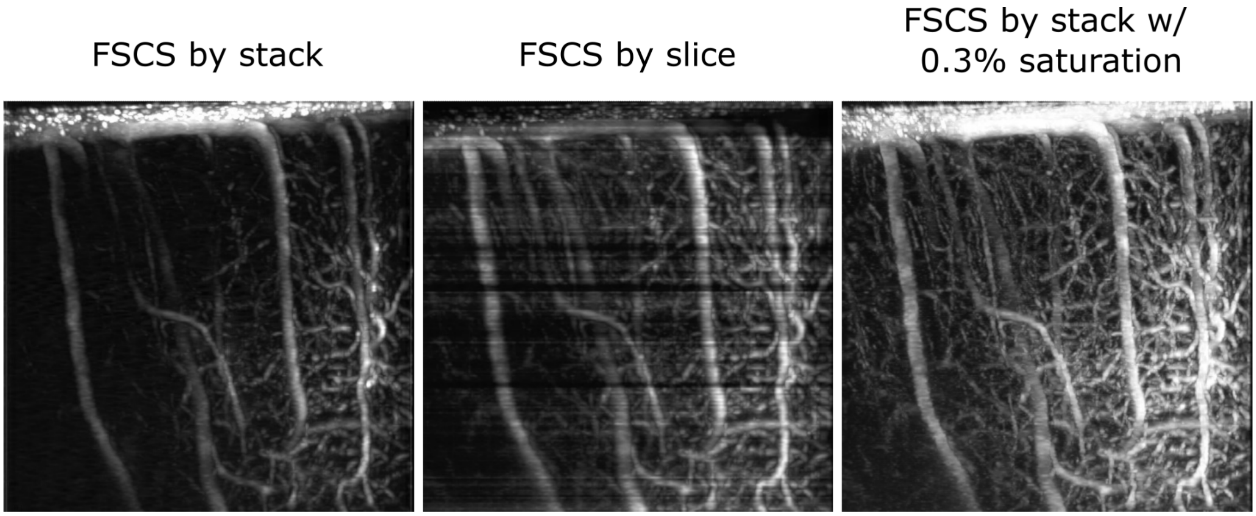


**S.Fig. 2. Test image results for models trained & tested with different normalization methods.** Sagittal projections of model output image stacks that underwent full scale contrast stretch (FSCS) with respect to the entire stack, per slice, or entire stack with 0.3% saturation prior to input to each respectively trained model as test images. FSCS was done with Fiji ImageJ normalization function[1].

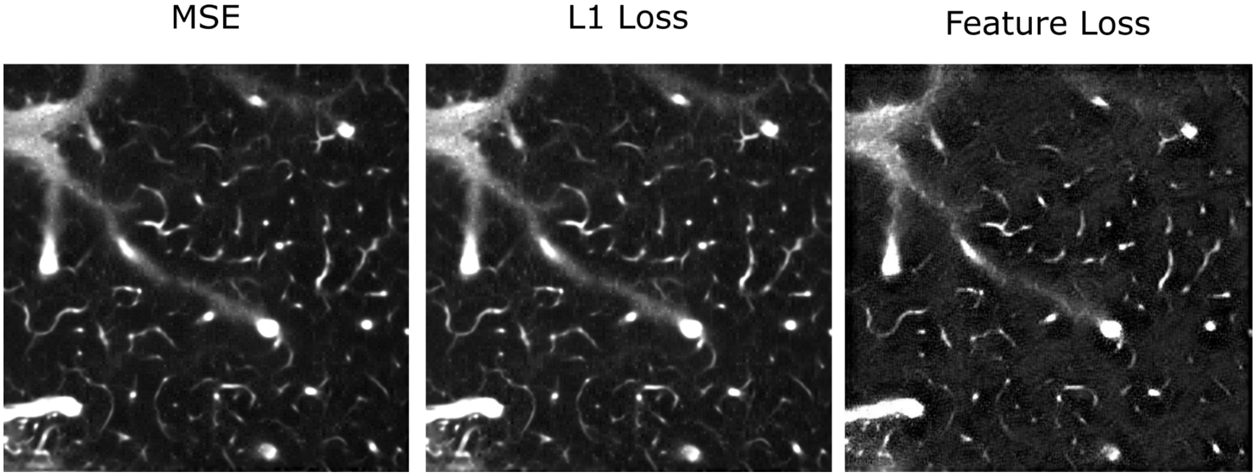


**S.Fig. 3. Test image results for models trained with different loss functions (single image slice).**
